# ForceFlowAb: physics-aware mixture-of-experts flow matching model for antibody CDRs design

**DOI:** 10.64898/2026.09.07.749983

**Authors:** Zonghui Li, Zexin Lv, Guijun Zhang

## Abstract

**Motivation:** Antibodies are a major class of therapeutic molecules, and their recognition of target antigens is largely mediated by complementarity-determining regions (CDRs), making antigen-conditioned CDR design a central problem in antibody engineering. Recent generative methods have enabled antigen-conditioned co-design of CDR sequences and structures, but their limited capacity to capture local interface heterogeneity and lack explicit energy-based guidance during sampling, which may result in unfavorable antibody-antigen interaction energies. Overcoming these limitations requires methods that better represent diverse interface environments via adaptive routing and incorporate physical guidance to steer sampling toward energetically favorable conformations.

**Results:** We present ForceFlowAb, a physics-aware mixture-of-experts flow-matching framework for antigen-conditioned CDR sequence–structure co-design. The framework models heterogeneous interface environments through specialized expert routing and applies differentiable force-field guidance during sampling to guide CDR generation toward energetically favorable conformations. For CDR-H3 design, ForceFlowAb achieved more favorable antibody–antigen interaction energies than FlowDesign and Diffab, with improvement rates (IMP) of 46.5% versus 35.0% and 35.5%, respectively. For simultaneous six-CDR design, ForceFlowAb also outperformed Diffab, with IMP values of 16% versus 9%. These results suggest complementary roles for interface-adaptive modeling and energy-based guidance, with the former capturing binding-mode diversity and the latter leveraging physical constraints to ensure biophysical feasibility.

**Availability and implementation:** The web server is freely available at http://zhanglab-bioinf.com/ForceFlowAb. The source code and implementation are available at https://github.com/iobio-zjut/ForceFlowAb.

**Supplementary information:** Supplementary data are available at Bioinformatics online.

## 1 Introduction

Antibodies are a major class of therapeutic molecules, with broad applications in infectious diseases, cancer, and immune-related disorders (Scott et al., 2012). However, experimental screening of the vast antibody sequence–structure space is prohibitively expensive, driving growing interest in computational design. A central goal of antibody design is to engineer antibody–antigen recognition, which is largely mediated by the six complementarity-determining region (CDR) loops (MacCallum et al., 1996; Sela-Culang et al., 2013). These loops exhibit diverse sequence and conformational preferences that collectively shape the physicochemical landscape of the binding interface, making rational CDR design central to engineering antibodies against predefined targets.

Traditional computational approaches to CDR design primarily relied on physics-based conformational sampling and energy optimization (Lippow et al., 2007). Methods such as RosettaAntibodyDesign (Adolf-Bryfogle et al., 2018) explored combinations of loop conformations and amino acid substitutions using all-atom energy functions (Alford et al., 2017). While physically interpretable, these approaches require extensive sampling and depend on the accuracy of the underlying energy functions, limiting their ability to efficiently explore the vast sequence–structure space of CDRs. To alleviate these limitations, language-modeling approaches were adapted to antibody sequences, leading to models such as AbLang (Olsen et al., 2022), AntiBERTy (Ruffolo et al.,2021), and IgLM (Shuai et al., 2023) that enable efficient and scalable CDR infilling and antibody sequence generation. Despite these advantages, their sequence-based representations do not explicitly capture three-dimensional CDR geometry, antigen context, or antibody–antigen interface complementarity. This limitation motivated the development of structure-aware approaches that explicitly incorporate geometric information into antibody design. Methods such as RefineGNN (Jin et al., 2022) and dyMEAN (Kong et al., 2023) introduced geometric graph representations and equivariant neural networks for antibody sequence–structure co-design. Compared with sequence-only models, these approaches jointly generate CDR sequences and conformations within the three-dimensional context of the antibody–antigen complex.

More recently, following their success across a broad range of generative modelling tasks, diffusion probabilistic models have been adapted to antibody design. By explicitly modelling the conditional joint distribution of CDRs sequences and structures, these models support diverse sampling under a given antigen context. Diffab (Luo et al., 2022) exemplified this direction by introducing an antigen-conditioned diffusion framework based on equivariant neural networks. AbDiffuser (Martinkus et al., 2023) extended this paradigm to full-atom antibody generation through an equivariant and physics-informed diffusion model, with subsequent in vitro validation. AbX (Zhu et al., 2024) further incorporated evolutionary, physical and geometric constraints into score-based diffusion to provide biologically and structurally informed priors. While, compared with conventional diffusion-based approaches to CDR co-design, flow matching formulates CDR generation as a transport process between flexible source and target distributions, thereby avoiding the need for specialized discrete diffusion processes to model categorical amino acid identities and reducing the computational cost of iterative denoising during large-scale sampling. FlowDesign (Wu et al., 2025) implemented this formulation, enabling informative prior selection, direct matching of discrete distributions and more efficient sampling.

Although generative models have substantially improved the efficiency of CDR co-design, dedicated constraints and guidance mechanisms are still required to ensure the quality of generated designs. First, antibody–antigen interfaces are locally heterogeneous, with different contact regions favouring distinct combinations of residue types, geometries and physicochemical interactions (MacCallum et al., 1996; Sela-Culang et al., 2013). While, existing co-design models typically use shared representations and parameters across diverse interface environments, which may limit their ability to capture region-specific geometric and physicochemical interaction patterns. Second, sequences and structures generated by deep learning models may not be physically feasible at the binding interface. Existing physical or geometric priors are often incorporated implicitly through model architectures or training objectives, leaving limited explicit energy-based guidance during sampling. Consequently, generated candidates may still exhibit steric clashes, unfavourable contacts or insufficient interfacial complementarity. Addressing these challenges requires an interface-adaptive co-design framework that captures heterogeneous local interaction patterns while using physical guidance to steer CDR conformations towards energetically favourable antibody–antigen interfaces.

Here, we present ForceFlowAb, a physics-aware mixture-of-experts flow-matching framework for antigen-conditioned CDR sequence–structure co-design. ForceFlowAb integrates three central design mechanisms. First, it formulates CDR co-design as a conditional flow-matching process that jointly transports sequence and structural states using contextual information from the antibody framework and target antigen. Second, an interface mixture-of-experts module routes local representations to specialized experts, enabling adaptive modelling of heterogeneous antibody– antigen interaction patterns. Third, a differentiable force-guidance mechanism steers generated CDR conformations towards energetically favourable antibody–antigen interfaces during sampling. Across CDR-H3 and simultaneous six-CDR co-design tasks, ForceFlowAb yielded favorable antibody–antigen interaction energies than representative generative baselines while maintaining competitive sequence recovery, structural accuracy and interface quality.

## 2 Materials and methods

### 2.1 Dataset preparation

Antibody-antigen complex structures were collected from the Structural Antibody Database (SAbDab) (Dunbar et al., 2014). We downloaded all antibody-antigen complexes available as of 18 October 2025, yielding an initial set of 10,056 structures. CDR regions were defined according to the Chothia numbering scheme (Chothia and Lesk, 1987; Chothia et al., 1989). The dataset was first cleaned by removing antibodies targeting non-protein antigens and structures with a resolution worse than 4 Å. To construct training, validation, and test splits while minimizing sequence leakage, all available CDR sequences from each antibody were concatenated and clustered using MMseqs2 (Steinegger and Söding, 2017) at a sequence identity threshold of 70%, following the clustering strategy used in RFantibody (Bennett et al., 2026). This procedure resulted in 3,377 non-redundant clusters. For fair comparison with FlowDesign, we adopted its test set of 20 antibody-antigen complexes. These complexes corresponded to 16 clusters in our clustered dataset, with no overlap with the training or validation sets. After excluding the clusters assigned to the test set, the remaining 3,361 clusters were divided into training and validation sets at a ratio of 9:1.

### 2.2 Task formulation

We derive a residue-level geometric representation for each antibody-antigen complex from its three-dimensional structure. For a complex with *N* residues, residue *i* is represented by its amino acid identity *s*_*i*_ ∈ {0, …, 19} encoded as a one-hot vector in ℝ^20^, its *Cα* coordinate *x*_*i*_ ∈ ℝ^3^, and a local backbone orientation frame *O*_*i*_ ∈ SO(3). The local frame is constructed from backbone atoms following the residue-frame parameterization used in Diffab (Luo et al., 2022). The target CDR segment is denoted as *R* = {*s*_*i*_, *x*_*i*_, *O*_*i*_ ∣ *i* = *l* + 1, …, *l* + *m*} where *m* is the length of the designed CDR. The remaining residues constitute the conditioning context *C* = {(*s*_*j*_, *x*_*j*_, *O*_*j*_) ∣ *j* ∈ {1, …, *N*} ∖ {*l* + 1, …, *l* + *m*}}. The goal of antigen-conditioned CDR design is to generate a CDR state *R*, conditioned on the fixed context *C*, such that the generated CDR exhibits plausible amino acid preferences, adopts a structurally valid loop conformation, and forms favorable interactions with the antigen.

ForceFlowAb formulates this task as a conditional transport problem (Lipman et al., 2023; Wu et al., 2025). Given an initial CDR state *T* = {*T*_*s*_, *T*_*x*_, *T*_*O*_} sampled from a prior distribution *p*_0_(. ∣ *C*), the model learns a conditional vector field that transports the prior distribution toward the empirical CDR distribution *p*_1_(*R* ∣ *C*). During training, a time point *t* ~ *U* (0,1) is sampled, and an interpolated CDR state *R*^*t*^ is constructed between the initial state *T* and the native state *R*. Amino acid identities and *Cα* coordinates are interpolated linearly:

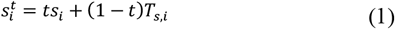

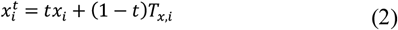

Backbone orientations are interpolated on SO(3). Specifically, the initial orientation *T*_*O,i*_ and native orientation *O*_*i*_ are converted into quaternion representations *T*_*q,i*_ and *q*_*i*_, respectively. Spherical linear interpolation (SLERP) is then applied (Shoemake, 1985):

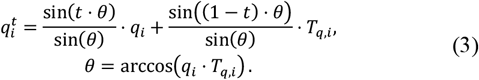

Then the interpolated quaternion is converted back to a rotation matrix. This construction provides a smooth interpolation path from *T*_*O,i*_ to *O*_*i*_ while keeping *O*^*t*^ ∈ SO(3). The resulting interpolated CDR state 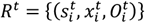, together with the fixed context *C* and time *t*, is used as input to the model.

### 2.3 Overview of ForceFlowAb

ForceFlowAb is a physics-aware mixture-of-experts flow-matching framework for antigen-conditioned antibody CDR design. Its architecture comprises five functional components: an Input Frame Embedder, a CDR Embedder, a MoE Module, a Decoder, and a Force Guidance module. The Input Frame Embedder encodes residue-level physicochemical features and pairwise geometric relationships derived from the antibody framework and antigen. The CDR Embedder combines the target CDR mask, antibody–antigen context, pairwise geometry, and flow-matching time step through iterative IPA and Transformer blocks to produce context-aware representations of the design region. The MoE Module further refines these representations through antigen cross-attention and expert routing, allowing specialized experts to model heterogeneous local interface environments. The Decoder maps the resulting representations to sequence, coordinate, and orientation drift fields, which iteratively update the generated CDR states along the flow trajectory. In the later phase of the sampling trajectory (specifically, steps 70–80), the Force Guidance module applies energy-derived coordinate guidance to the generated CDR Cα atoms, reducing unfavorable interactions and promoting physically plausible antibody–antigen interfaces. The overall pipeline of ForceFlowAb is illustrated in Fig. 1.

**Figure 1.**
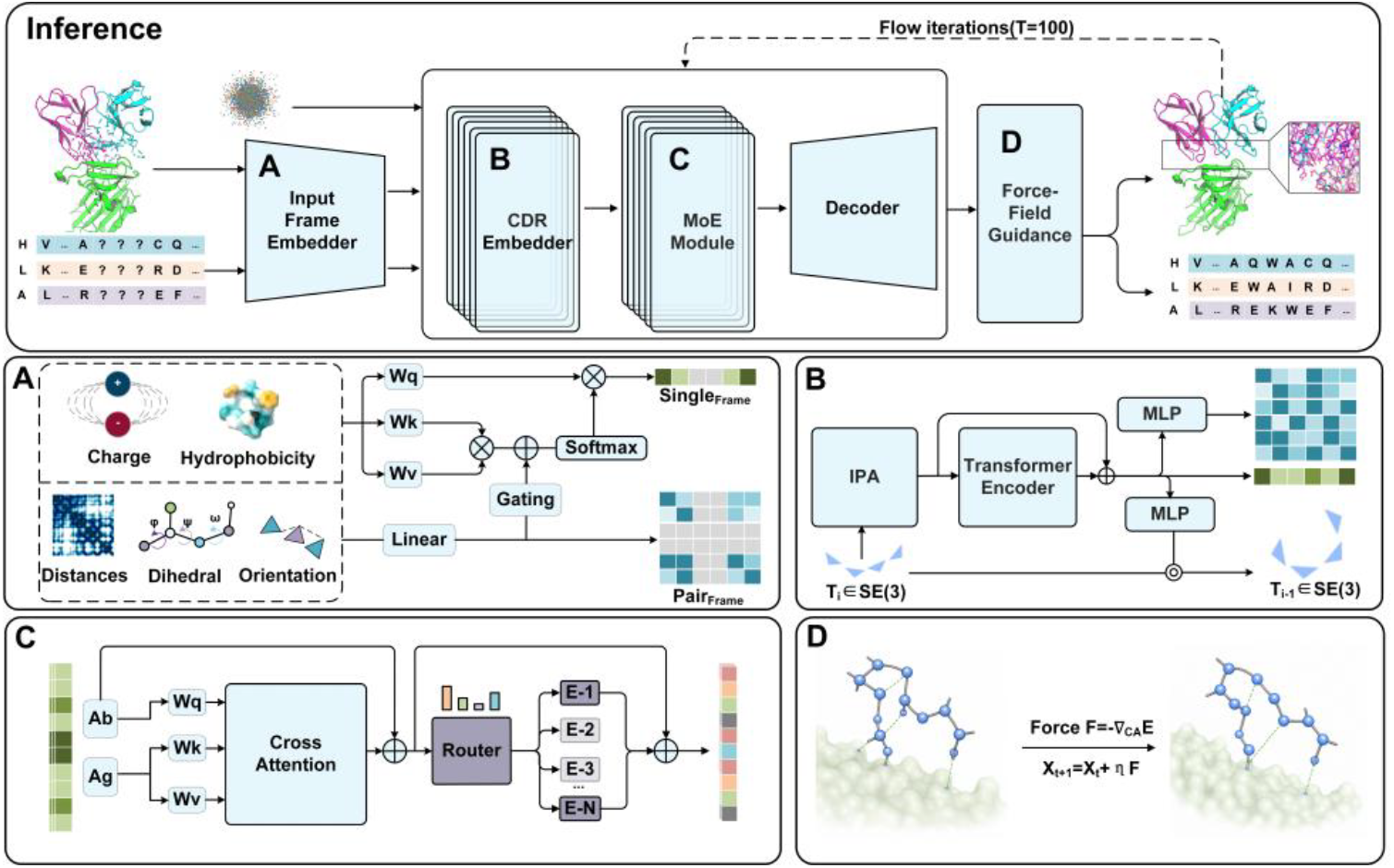
Overall pipeline of ForceFlowAb. The Input Frame Embedder encodes antibody–antigen residue and pairwise features. The CDR Embedder then integrates these features with the flow-matching time step through IPA and Transformer blocks, yielding time-dependent representations at the masked CDR positions. The MoE Module refines interface representations through antigen cross-attention and expert routing. The Decoder predicts sequence, coordinate, and orientation drift fields to iteratively update the CDR states, while energy-derived forces guide the generated CDR Cα coordinates toward physically favorable interface conformations.

#### 2.3.1 Input Frame Embedder

Each antibody–antigen complex is represented by residue-level and residue-pair features that jointly characterize the local environment of the designed CDRs. Residue-level features include amino acid identity, residue-local heavy-atom coordinates, backbone dihedral angles, chain type, hydropathy, and charge. Residue-pair features include amino-acid pair identity, a same-chain indicator, relative sequential positions, pairwise heavy-atom distances, and relative backbone orientations. Detailed definitions and dimensionalities of these features are provided in Supplementary Table S1. For fixed framework and antigen residues, these conditioning features are derived from the input complex and remain unchanged throughout generation. To prevent information leakage, native amino acid identities at masked CDR positions are replaced with an unknown-residue token, while their coordinate and dihedral features, together with the associated pairwise distance and orientation features, are masked. Only chain information, same-chain indicators, relative sequential positions, and unknown-residue-derived features are retained. The resulting features are then processed by an embedding network to generate the frame single and pair representations.

#### 2.3.2 CDR Embedder

Based on the frame single and pair representations produced by the Input Frame Embedder, the CDR Embedder propagates contextual information from the surrounding antibody framework and antigen residues to the masked CDR positions. The flow-matching time step t is embedded and added to the target CDR representations to condition the module on the current stage of generation. The resulting representations are iteratively refined using IPA and Transformer blocks (Jumper et al., 2021; Vaswani et al., 2017). IPA captures geometry-aware interactions in residue-local frames, whereas the Transformer models long-range contextual dependencies across the antibody–antigen complex. Through these iterative updates, the masked CDR positions acquire time-conditioned and context-aware representations that encode constraints from the antibody framework, loop continuity, the local antibody–antigen interface environment, and physicochemical compatibility.

#### 2.3.3 Mixture-of-experts module

Antibody-antigen interfaces are locally heterogeneous, and different interface regions may benefit from different representation transformations. The mixture-of-experts (Shazeer et al., 2017) module provides a flexible mechanism for this setting by learning expert-specific transformations and adaptively combining them through residue-dependent gating weights to refine antibody-side representations.

The MoE module first updates the antibody-side representations through cross-attention with the antigen-side representations. For each antibody-antigen complex, residue representations are partitioned into antibody-side representations *H*_Ab_ and antigen-side representations *H*_Ag_ according to chain-type labels. The antigen representations serve as keys and values, while the antibody representations serve as queries. For an antibody position *i* the hidden representation *h*_*i*_ is updated as:

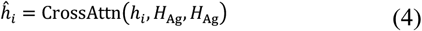

The updated representation 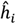is then passed to the MoE. For each antibody position *i*, a gating network maps 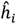 to a routing weight vector over all experts:

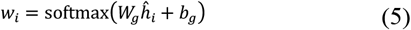

where *W*_*g*_ and *b*_*g*_ are learnable gating parameters, and *w*_*i,e*_ denotes the routing weight assigned to expert e for position *i*.

Each expert is implemented as an independent feed-forward network (FFN), which performs an expert-specific transformation on the representation:

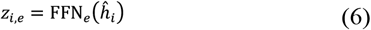

where FFN_*e*_ denotes the *e*-th expert network. The expert outputs are aggregated according to the routing weights:

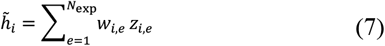

where *N*_exp_ is the number of experts.

After the MoE module, the resulting representation 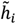is fed into three independent MLP heads to predict the sequence, coordinate, and orientation drift fields, denoted by *F*_*θ,i*_, *G*_*θ,i*_ and *H*_*θ,i*_ respectively, for each target CDR position.

#### 2.3.4 Force-field-guided sampling

MadraX is a knowledge-based, fully differentiable protein force field implemented as a PyTorch module (Orlando et al., 2024). Inspired by the FoldX energy function (Guerois et al., 2002; Schymkowitz et al., 2005), MadraX expresses the conformational energy of proteins and protein complexes as a sum of differentiable physicochemical terms. In ForceFlowAb, we used a reduced MadraX-based guidance energy during sampling, retaining selected structure-preserving terms rather than summing all available MadraX energy terms. This reduced formulation was adopted to improve gradient stability by omitting energy terms that are highly sensitive to detailed side-chain conformations. The retained terms comprised backbone hydrogen-bond energy, backbone entropy, and the peptide-bond violation penalty. During flow sampling steps 70–80, the guidance energy was evaluated on the intermediate antibody–antigen complex formed by the current generated CDR state and the fixed surrounding context. Its gradient was computed only with respect to the generated CDR Cα coordinates, while the antibody framework and antigen coordinates were kept fixed. Specifically, the physical guidance force was calculated as:

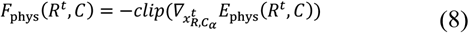

where *E*_phys_(*R*^*t*^, *C*) is the reduced MadraX guidance energy and 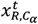denotes the generated CDR *Cα* coordinates at time *t*. The clipped physical gradient was scaled by *G*_*θ*_ and integrated into the coordinate update.

### 2.4 Loss Function

The training objective comprises three components: vector-field matching losses, a CDR hydropathy preference loss, and an auxiliary MoE balancing loss.

The vector-field matching loss supervises the predicted sequence, coordinate, and orientation drift fields for the target CDR residues and is defined as:

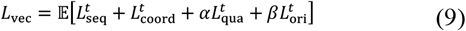

where the expectation is taken over training complexes, prior samples, and time steps. *α* and *β* are the weighting coefficients for the quaternion and orientation losses, respectively, and their values are provided in the Supplementary Note 1. The individual loss terms are given by

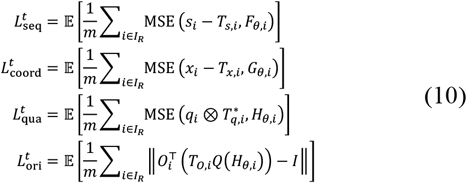

where *I*_*R*_ denotes the set of target CDR residues, *m* = |*I*_*R*_|, *q*_*i*_ and *T*_*q,i*_ are the native and initial quaternions, 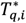 denotes the quaternion conjugate, ⊗ denotes quaternion multiplication, and *Q*(⋅) converts the predicted quaternion representation into a rotation matrix.

To regularize the physicochemical properties of generated CDR sequences, we introduce a hydropathy loss based on the Kyte-Doolittle hydropathy scale (Kyte and Doolittle, 1982). Let *r*_*a*_ denote the normalized hydropathy value of amino acid type a.

For residue *i*, the predicted and native hydropathy values are computed as 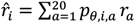, and 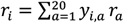 respectively. where *p*_*θ,i,a*_ is the predicted amino acid probability and *y*_*i,a*_ is the native one-hot label. The hydropathy loss is defined as

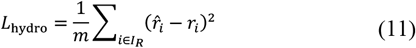

To prevent routing collapse in the MoE module, we introduced an auxiliary load-balancing loss. Let *p*_*b,i,e*_ denote the softmax routing probability assigned to expert *e* for residue *i* in sample *b*, and let *z*_*b,i,e*_ be a binary indicator of whether expert *e* is selected among the top-*K* routed experts. For each sample, we define two per-expert routing statistics, the normalized expert assignment frequency 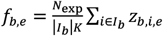 and the average routing probability 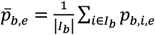.

The MoE balancing loss is then computed as

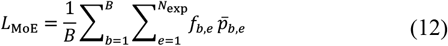

where *N*_exp_ is the number of routed experts, *K* is the number of experts selected per token, *I*_*b*_ denotes the valid residue tokens in sample *b*.

The final training objective is:

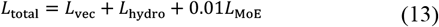

## 3 Results

### 3.1 Evaluation settings and baselines

We compared ForceFlowAb with two representative generative antibody design methods, Diffab and FlowDesign. AbX and AbDiffuser were also considered but excluded because neither could be deployed under our unified evaluation protocol. The official AbX release provides pretrained checkpoints and inference and evaluation utilities but no training pipeline, whereas no official implementation of AbDiffuser was available at the time of our experiments. For a fair comparison, Diffab and FlowDesign were retrained following their original training pipelines using the same training and validation sets described in Section 2.1. For each test complex, each method sampled 100 candidate structures. After inference, side chains were constructed using the side-chain packing function in Rosetta, followed by relaxation to reduce interatomic clashes. The relaxed candidates were ranked by Rosetta interface energy, and the top 10 candidates were retained for evaluation. Rosetta interface energy was calculated using the InterfaceAnalyzerMover in PyRosetta with the REF2015 weight function, representing a weighted sum of Rosetta energy terms at the antibody–antigen interface (Chaudhury et al., 2010; Alford et al., 2017). All methods were evaluated on the same held-out test set using an identical side-chain packing, relaxation and candidate-selection procedure.

### 3.2 Metrics

We evaluated generated CDRs at the sequence, structural, and interface levels. At the sequence level, amino acid recovery rate (AAR) was used to quantify agreement with the native CDR sequence. At the structural level, Cα root-mean-square deviation (Cα RMSD) was used to measure the accuracy of the designed CDR conformations. At the interface level, Rosetta interface energy, improvement rate (IMP), and DockQ (Basu and Wallner, 2016) were used to assess antibody–antigen interface quality. Lower Rosetta interface energy indicates more favorable antibody–antigen interactions, and IMP was defined as the proportion of generated candidates with lower interface energy than the corresponding reference complex. In addition, AlphaFold 3 was used as an independent assessment of sequence–structure compatibility, with designed-CDR pLDDT reflecting local structural confidence and ipTM reflecting predicted antibody–antigen interface confidence (Mariani et al., 2013; Abramson et al., 2024).

### 3.3 Antigen-binding CDR-H3 design

We evaluated all methods on antigen-conditioned CDR-H3 design. As shown in Fig. 2a, ForceFlowAb tended to produce lower ΔΔG values calculated from Rosetta interface energy than FlowDesign and Diffab, suggesting more favorable predicted antibody–antigen interface energetics. ForceFlowAb also showed a higher improvement rate (IMP; Fig. 2d) and higher DockQ scores than the baselines (Fig. 2c), indicating improved interface quality. For CDR-H3 structural accuracy, ForceFlowAb showed Cα RMSD comparable to FlowDesign and Diffab, with no significant differences among the three methods (Fig. 2b). In terms of sequence recovery, ForceFlowAb achieved the highest AAR, with a value comparable to FlowDesign and higher than Diffab (Fig. 2e). Detailed numerical results are provided in Supplementary Table S2. Together, these results suggest that interface-adaptive modelling and energy-based guidance jointly contribute to improved antibody–antigen interaction energies and interface quality.

**Figure 2.**
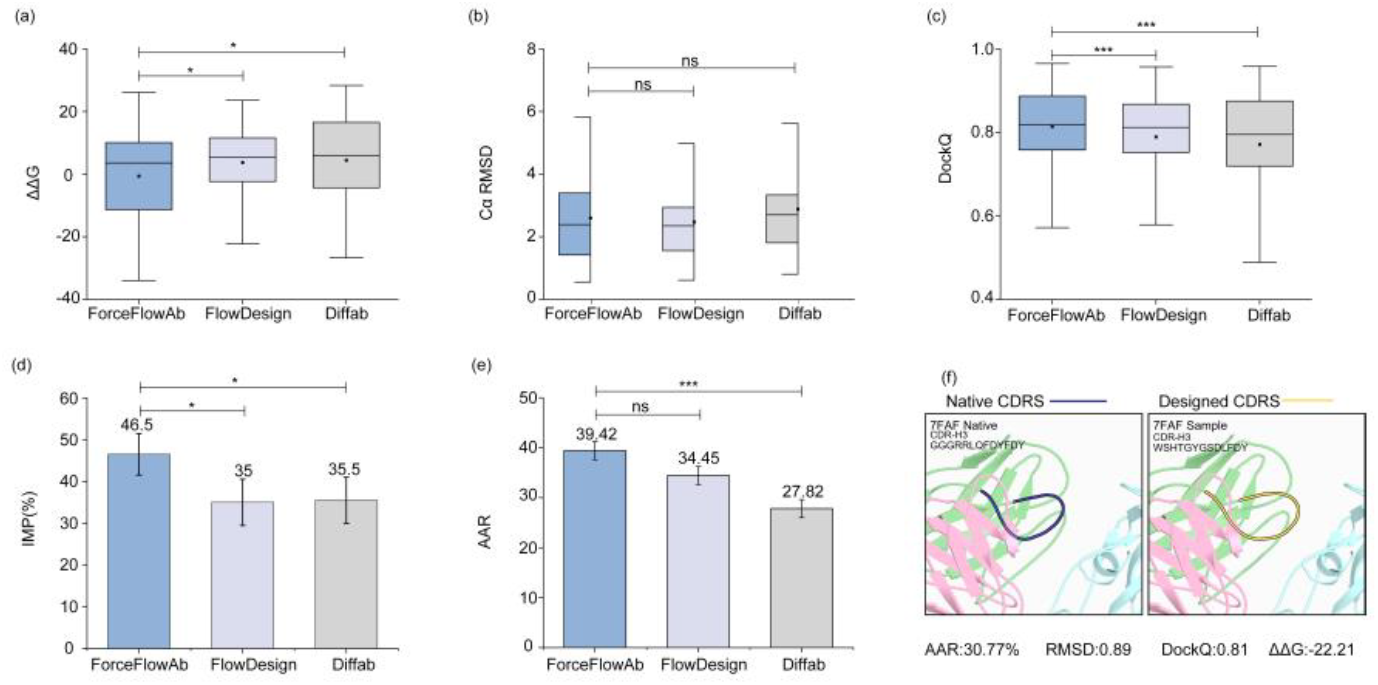
Evaluation of antigen-conditioned CDR-H3 design. (a–c) Comparison of ΔΔG values calculated from Rosetta interface energy, CDR-H3 Cα RMSD, and DockQ among ForceFlowAb, FlowDesign, and Diffab. Square markers indicate mean values. (d,e) Improvement rate (IMP) and amino acid recovery rate (AAR). (f) Representative SARS-CoV-2 antibody-antigen complex (PDB: 7FAF), showing the native and ForceFlowAb-generated CDR-H3 loops together with the corresponding sequence and evaluation metrics. Error bars indicate 90% confidence intervals. ns, not significant; *P < 0.05; ***P < 0.001.

### 3.4 Antigen-binding six-CDR design

We next evaluated ForceFlowAb on simultaneous six-CDR design, in which H1, H2, H3, L1, L2, and L3 were jointly masked and generated while the antibody framework and antigen were kept fixed. In this setting, ForceFlowAb was compared with Diffab. FlowDesign was not included because its public implementation does not support simultaneous multi-CDR design under the same evaluation protocol. As shown in Fig. 3a, ForceFlowAb produced lower Rosetta interface energy than Diffab in the six-CDR design setting. ForceFlowAb also showed a higher improvement rate than Diffab (16% versus 9%; Fig. 3b), suggesting more favorable predicted antibody–antigen interface energetics. Additional analyses of Cα RMSD, DockQ, and AAR showed that ForceFlowAb maintained generally favorable performance across the six CDR loops (Supplementary Fig. S1). Detailed numerical results are provided in Supplementary Table S3 and Supplementary Table S4. These results indicate that ForceFlowAb can be extended beyond CDR-H3 design to the more challenging setting of simultaneous multi-loop CDR co-design. Representative six-CDR design examples are provided in Supplementary Fig. S2.

**Figure 3.**
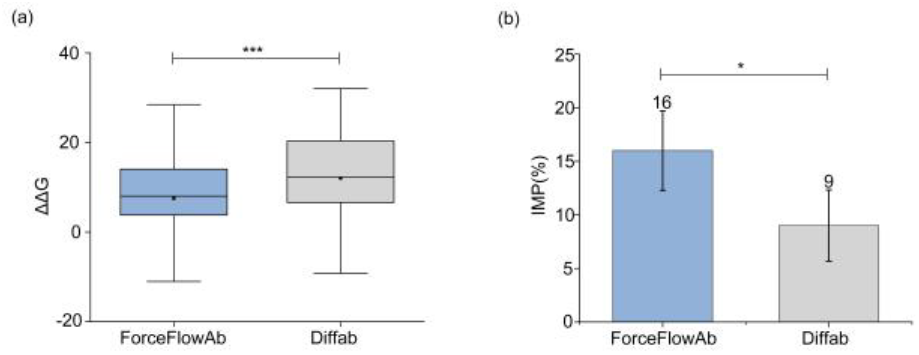
Evaluation of simultaneous six-CDR design. (a) Distribution of ΔΔG values calculated from Rosetta interface energy. Square markers indicate mean values. (b) Improvement rate (IMP) based on ΔΔG. Error bars indicate 90% confidence intervals. *P < 0.05; ***P < 0.001.

### 3.5 AlphaFold 3-based plausibility assessment of generated CDR sequences

Following RFantibody’s retrospective analysis, which suggested that the AF3 ipTM score may be useful for filtering antibody designs (Bennett et al., 2026), we used AlphaFold 3 as an additional computational assessment of the sequence–structure compatibility of generated CDRs. For the main analysis, we focused on simultaneous six-CDR design and compared ForceFlowAb with Diffab. For each test complex, 100 candidate sequences were generated, each of which was independently predicted by AF3, and the top 10 candidates ranked by ipTM were retained for analysis. The predicted complexes were evaluated using ipTM and designed-CDR pLDDT, reflecting predicted interface confidence and local structural confidence, respectively. Following RFantibody, ipTM = 0.6 was used as an interface-confidence reference (Bennett et al., 2026), while CDR-pLDDT = 70 was used as a local-confidence reference based on the AlphaFold confidence scale (Jumper et al., 2021; Abramson et al., 2024). We further defined the joint confidence pass rate as the proportion of retained candidates exceeding both thresholds. As shown in Fig. 4, ForceFlowAb achieved a higher pass rate than Diffab (50% versus 42%), suggesting improved compatibility between the designed CDR sequences and plausible antibody–antigen complex conformations. AF3-based assessment for CDR-H3 design is provided in Supplementary Fig. S3, while selected six-CDR design examples are shown in Supplementary Fig. S4.

**Figure 4.**
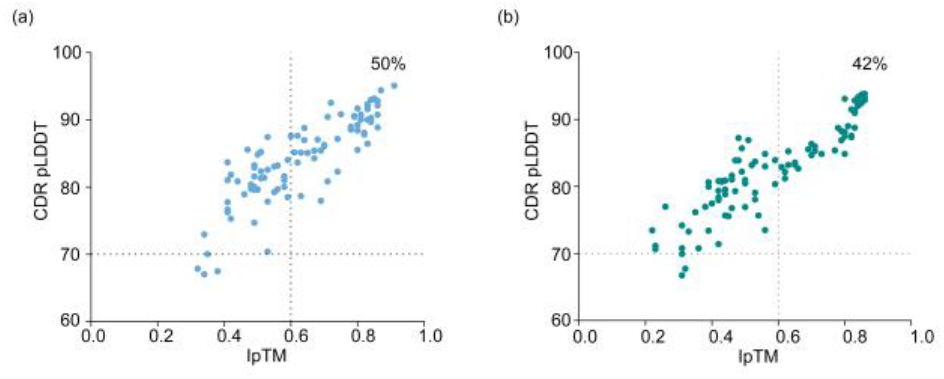
AlphaFold 3-based assessment of generated six-CDR sequences. Distributions of ipTM and designed-CDR pLDDT are shown for ForceFlowAb (a) and Diffab (b). Dashed lines indicate the reference values of ipTM = 0.6 and CDR-pLDDT = 70. Percentages indicate the joint confidence pass rate, defined as the proportion of retained candidates exceeding both reference values.

### 3.6 Expert selection analysis

To analyze expert selection, we extracted the routing outputs from the final MoE layer. For each CDR token, the two experts with the highest routing weights were recorded. For CDR region CDR_*i*_ and Expert *e*, top-2 routing enrichment was defined as

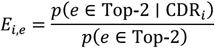

where *i* indexes the six CDR regions, *p*(*e* ∈ Top-2 ∣ CDR_*i*_) is the frequency with which Expert *e* was selected among the top two experts for tokens in CDR_*i*_, and *p*(*e* ∈ Top-2) is its overall top-2 selection frequency across all analyzed CDR tokens. Values greater than 1 indicate enrichment relative to the overall expert usage. As shown in Fig. 5, expert preferences varied across CDR regions and between antibody–antigen complexes, indicating that expert selection depends on the input CDR environment rather than following a fixed correspondence between experts and CDR identities. Detailed numerical results are provided in Supplementary Fig. S5.

**Figure 5.**
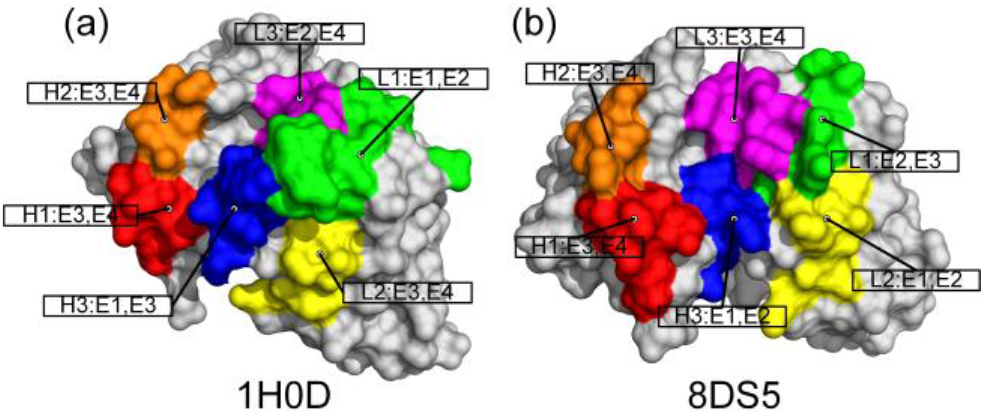
Expert selection patterns across CDR loops in two representative antibody–antigen complexes. Expert selection patterns are shown for (a) PDB 1H0D and (b) PDB 8DS5. The corresponding antibody surface representation shows the six CDR loops in different colors, with annotations indicating their two most enriched experts. Antibody framework regions (non-CDR regions) are shown in grey.

### 3.7 Ablation

To assess the contributions of individual components in ForceFlowAb, we performed ablation studies under the simultaneous six-CDR design setting by comparing the full model with three variants: removing force-guided sampling, removing the interface MoE module, and removing both components. As shown in Fig. 6 these analyses demonstrated that the interface MoE module mainly contributes to interface-aware sequence and structural modeling, whereas force-guided sampling provides additional energetic refinement during generation. The combined removal of both components resulted in a consistent degradation of model performance, supporting their complementary roles in improving antibody–antigen interface design. The effects of different ablation settings on AAR are shown in Supplementary Fig. S6, while additional parameter analyses of the MoE module and force-guided sampling strategy are presented in Supplementary Figs S7 and S8, respectively.

**Figure 6.**
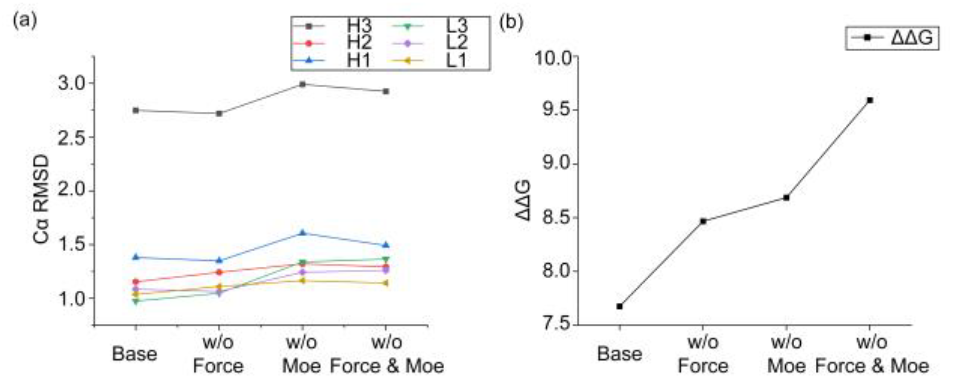
Ablation analysis under simultaneous six-CDR design. The full ForceFlowAb model is compared with variants removing force guidance, the interface MoE module, or both. (a) Cα RMSD of designed CDR loops. (b) ΔΔG values calculated from Rosetta interface energy, where lower values indicate more favorable interface energetics.

## 4 Conclusion

In this work, we presented ForceFlowAb, a physics-aware mixture-of-experts flow-matching framework for antigen-conditioned antibody CDR sequence-structure co-design. ForceFlowAb combines conditional flow matching, interface-adaptive expert routing and force-derived coordinate guidance to model heterogeneous antibody-antigen interfaces while improving physical plausibility during generation. Across CDR-H3 and simultaneous six-CDR design tasks, ForceFlowAb achieved favorable performance compared with representative diffusion and flow-matching-based baselines. It improved interface-related metrics, including Rosetta energy, IMP and DockQ, while maintaining competitive sequence recovery and CDR Cα RMSD. Independent AlphaFold 3 evaluation further supported the structural compatibility of the generated sequences.

From a practical perspective, future work will focus on experimental validation and further methodological improvement. In particular, wet-lab validation is needed to determine whether the generated antibodies can be experimentally produced and bind effectively to the target antigen. In addition, improving the efficiency of physical guidance and extending ForceFlowAb to less reliable or predicted complex structures may further enhance its practical utility in antibody discovery.

## Data availability

The source code and processed data underlying this article are publicly available in the ForceFlowAb GitHub repository at https://github.com/iobio-zjut/ForceFlowAb and are archived on Zenodo at https://doi.org/10.5281/zenodo.21645253.

## Supplementary

**Supplementary Table S1.** Input features to the model.

| Feature | Raw shape | Description |
| --- | --- | --- |
| Amino acid identity | [N] | Categorical identity of each residue. The vocabulary contains the 20 standard amino acids and an unknown-residue token used at masked CDR positions. |
| Local heavy-atom coordinates | [N, A, 3] | Cartesian coordinates of the heavy atoms in each residue, transformed into a residue-local frame defined by the N, Cα, and C atoms. By default, A = 15. |
| Backbone dihedral angles | [N, 3] | Backbone torsion angles $\omega$, $\phi$, and $\psi$ for each residue. |
| Chain type | [N] | Categorical label indicating whether each residue belongs to the antibody heavy chain, antibody light chain, or antigen. |
| Hydropathy | [N, 1] | Normalized Kyte–Doolittle hydropathy value associated with each residue type. |
| Charge | [N, 3] | One-hot representation of three residue charge categories: negative, neutral, and positive. |
| Amino-acid pair identity | [N, N] | Joint categorical identity of the amino-acid types at residues i and j. |
| Same-chain indicator | [N, N, 1] | Binary indicator specifying whether residues i and j belong to the same protein chain. |
| Relative sequential position | [N, N, 1] | Signed sequence-index difference between residues i and j, clipped to [-32, 32]. This feature is masked for cross-chain residue pairs. |
| Pairwise heavy-atom distances | [N, N, A<sup>2</sup>] | Euclidean distances between all valid heavy-atom pairs of each residue pair. |
| Relative backbone orientations | [N, N, 2] | Two inter-residue dihedral angles describing the relative backbone orientation of each residue pair, calculated from their N, Cα, and C atoms. |

*Note:* $N$ denotes the number of residues in antibody–antigen structural context, and $A$ denotes the number of heavy-atom slots per residue.

**Supplementary Table S2.**
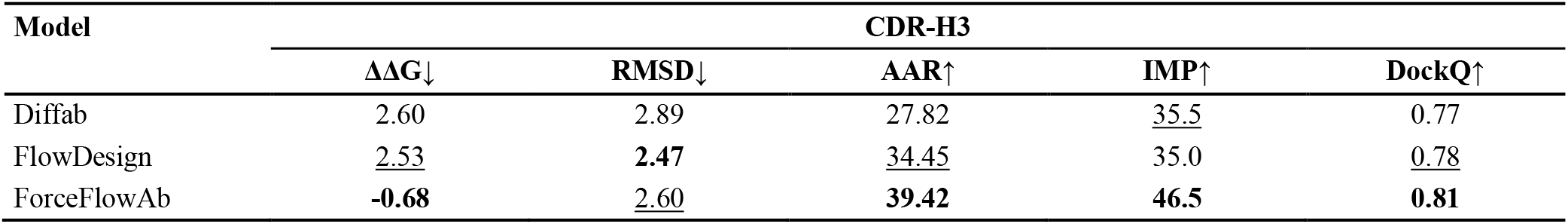
Mean performance metrics of CDR-H3 design.

**Supplementary Table S3.** Mean performance metrics of Six-CDR design.

| Model | Six-CDR |  |  |
| --- | --- | --- | --- |
| | $\Delta\Delta G\downarrow$ | IMP↑ | DockQ↑ |
| Diffab | <u>12.03</u> | <u>9</u> | <u>0.64</u> |
| ForceFlowAb | **7.67** | **16** | **0.73** |

**Supplementary Table S4.** Mean performance metrics of Six-CDR design.

| Model | Six-CDR |  |  |  |  |  |  |  |  |  |  |  |
| --- | --- | --- | --- | --- | --- | --- | --- | --- | --- | --- | --- | --- |
|  | H1 |  | H2 |  | H3 |  | L1 |  | L2 |  | L3 |  |
| | RMSD↓ | AAR↑ | RMSD↓ | AAR↑ | RMSD↓ | AAR↑ | RMSD↓ | AAR↑ | RMSD↓ | AAR↑ | RMSD↓ | AAR↑ |
| Diffab | **1.34** | <u>65.57</u> | <u>1.20</u> | <u>37.64</u> | <u>3.04</u> | <u>26.22</u> | <u>1.05</u> | <u>56.39</u> | **0.95** | <u>54.49</u> | <u>1.45</u> | <u>48.84</u> |
| ForceFlowAb | <u>1.37</u> | **82.71** | **1.15** | **69.78** | **2.74** | **38.34** | **1.03** | **86.66** | <u>1.08</u> | **88.92** | **0.97** | **79.02** |

**Figure S1.**
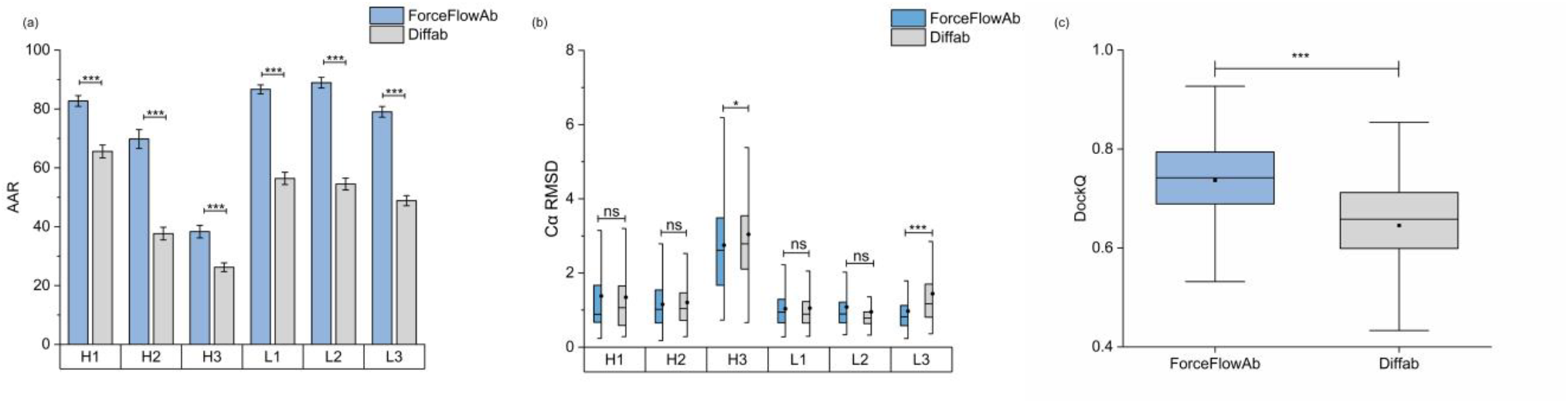
Additional evaluation of simultaneous six-CDR design. (a) Loop-wise amino-acid recovery rate (AAR) for H1, H2, H3, L1, L2, and L3. (b) Loop-wise CDR Cα RMSD distributions. (c) DockQ distributions of generated antibody-antigen complexes. ForceFlowAb is compared with Diffab under the same six-CDR design setting. Error bars indicate 90% confidence intervals. ns, not significant; *P < 0.05; ***P < 0.001. Detailed numerical results are provided in Supplementary Table S3 and Supplementary Table S4.

**Figure S2.**
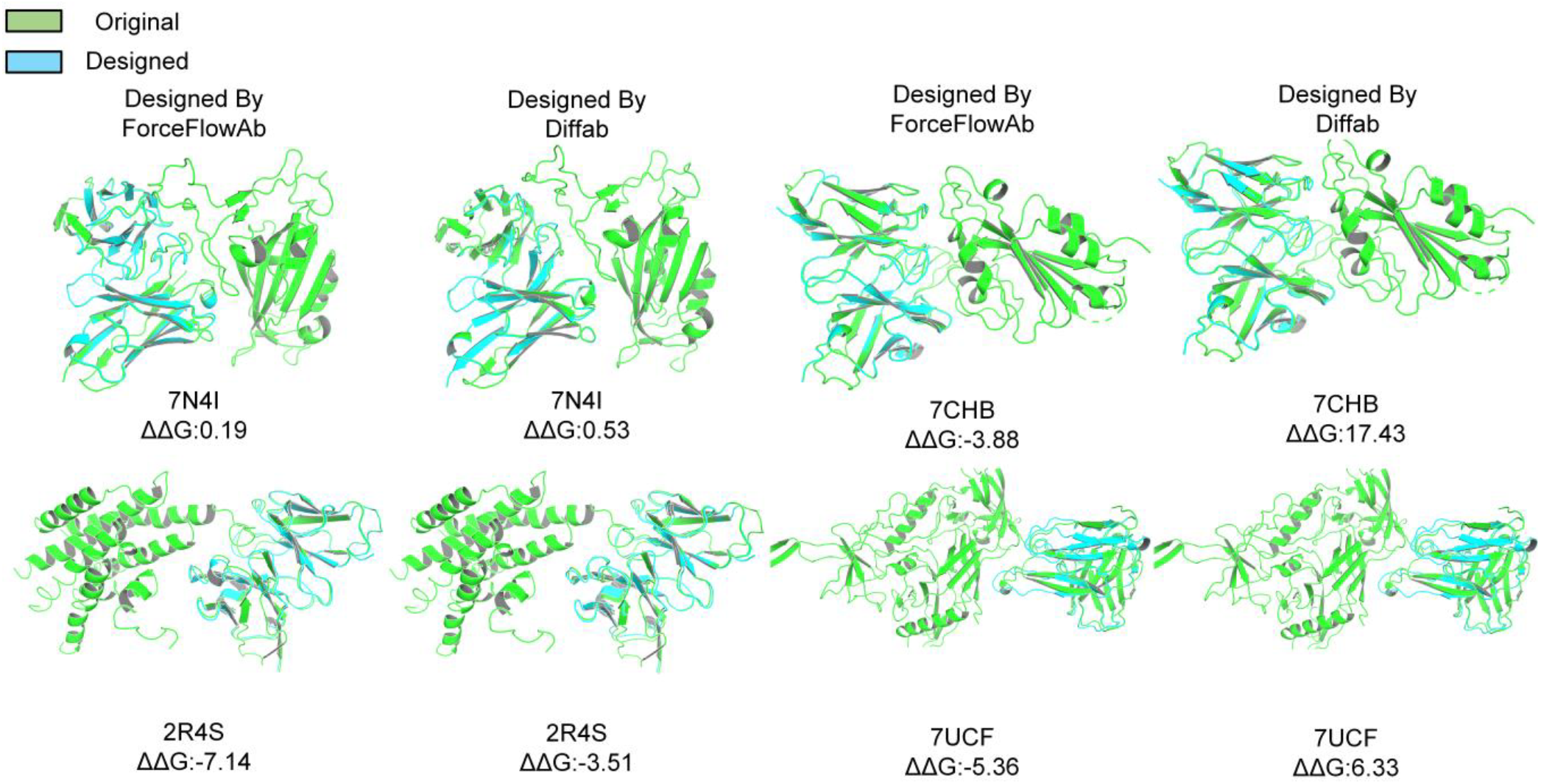
Representative examples of simultaneous six-CDR design. Original structures and generated structures are shown for ForceFlowAb and Diffab on four representative test complexes. Rosetta interface energy differences (ΔΔG) are reported below each model.

**Figure S3.**
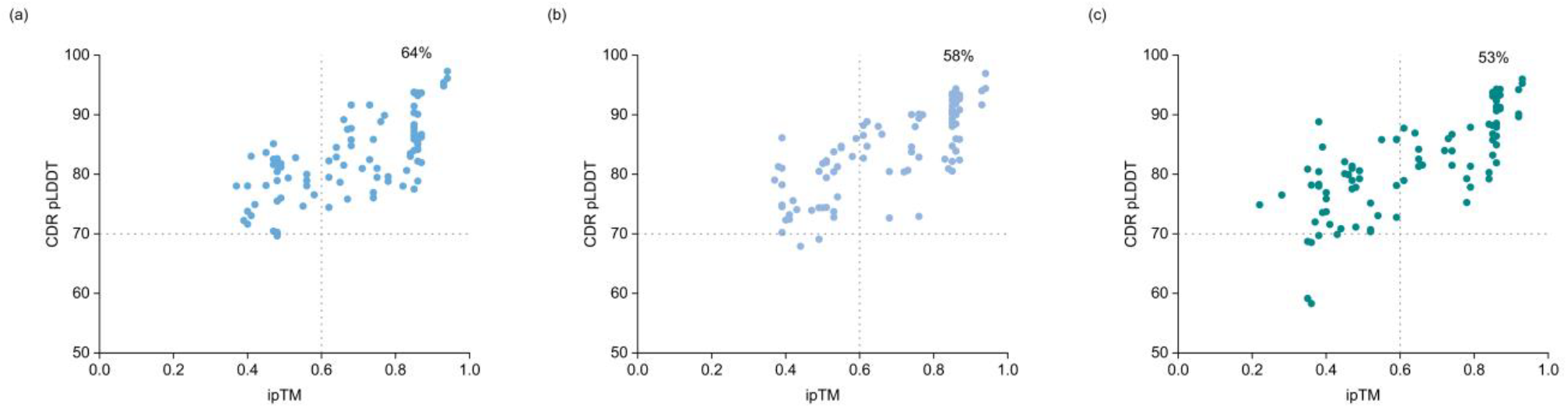
AlphaFold 3-based assessment of generated CDR-H3 sequences. Distributions of ipTM and designed-CDR pLDDT are shown for ForceFlowAb (a), FlowDesign (b), and Diffab (c). Dashed lines indicate the reference values of ipTM = 0.6 and CDR-pLDDT = 70. Percentages indicate the proportion of retained candidates exceeding both reference values.

**Figure S4.**
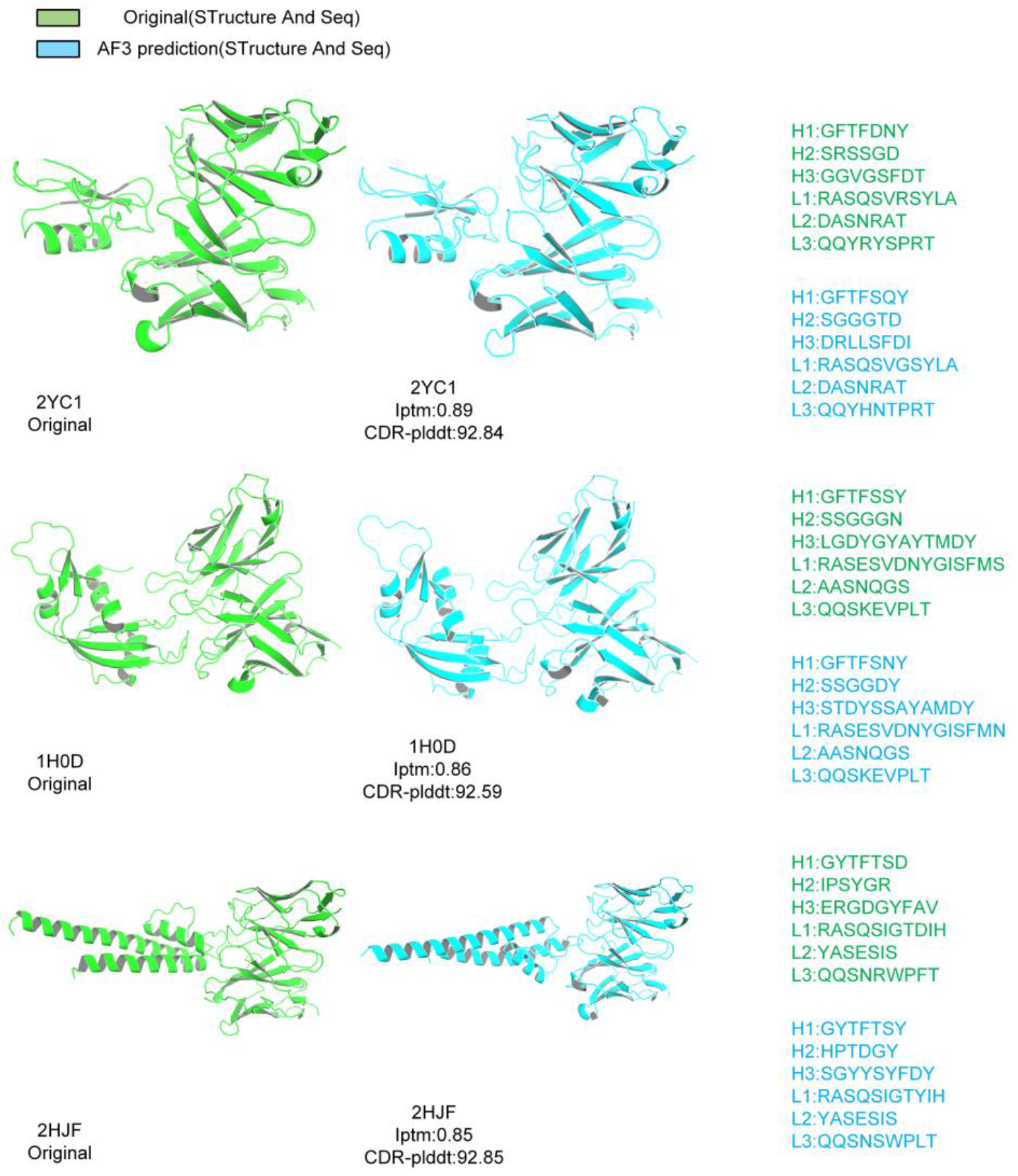
Representative comparison between native structures and AlphaFold 3 predictions of ForceFlowAb-designed sequences. Native structures and CDR sequences are shown in green, and AF3-predicted structures with designed CDR sequences are shown in blue. ipTM and designed-CDR pLDDT values are reported for each case

**Figure S5.**
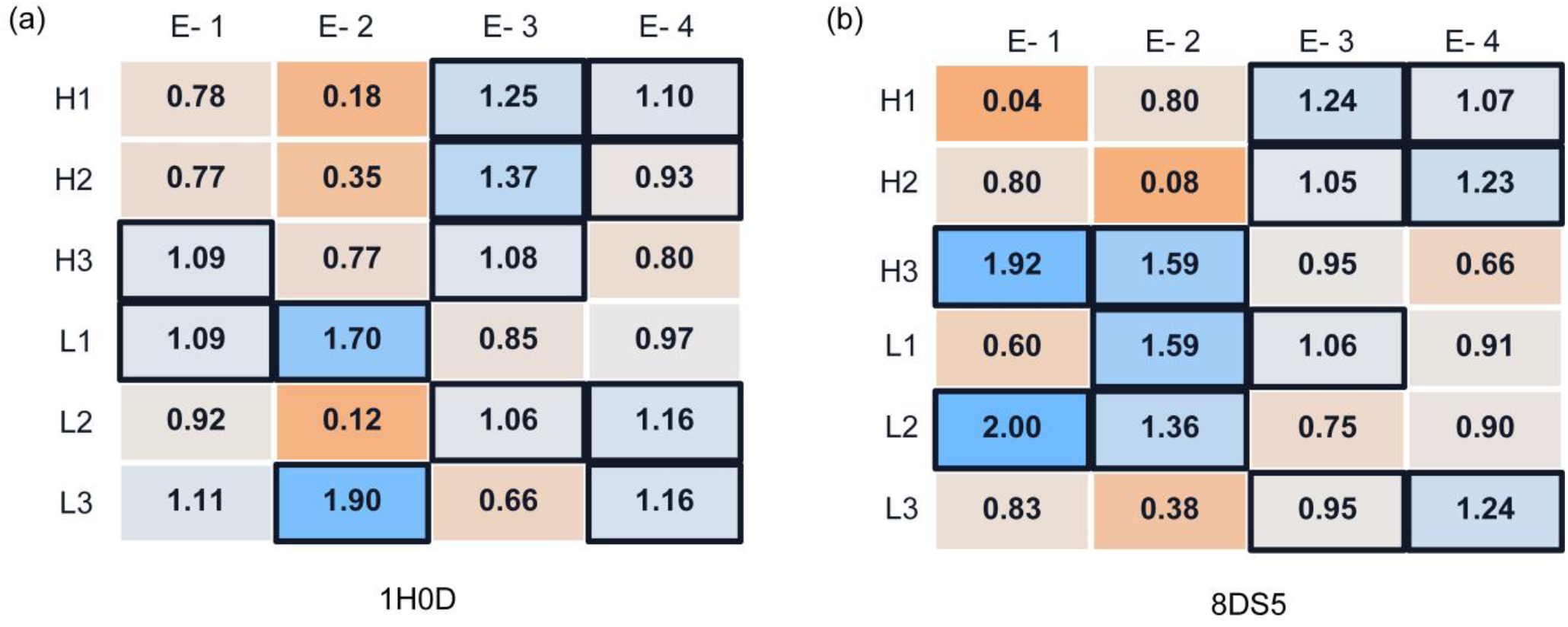
Expert selection patterns are shown for (a) PDB 1H0D and (b) PDB 8DS5. In each panel, the heatmap shows the top-2 routing enrichment of the four routed experts across the six CDR regions, with outlined cells indicating the two most enriched experts in each region.

**Figure S6.**
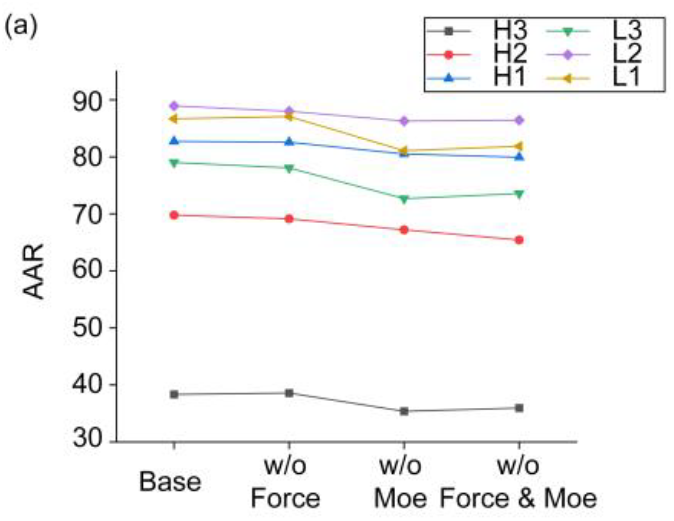
Ablation analysis under simultaneous six-CDR design. The full ForceFlowAb model is compared with variants removing force guidance, the interface MoE module, or both. (a) Amino acid recovery rate (AAR).

**Figure S7.**
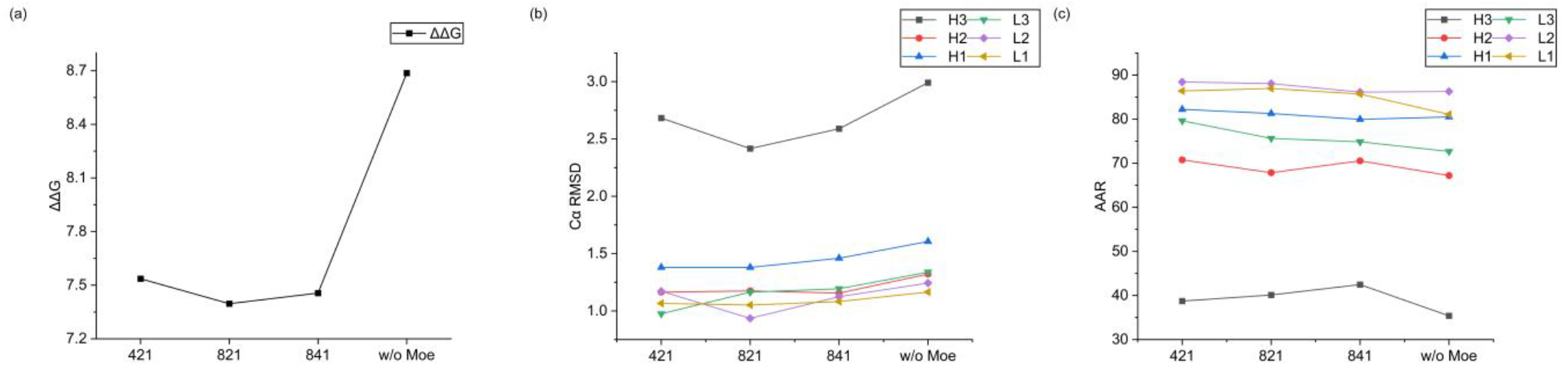
Ablation analysis of the interface MoE module under the simultaneous six-CDR design setting. Different MoE configurations are compared with the variant without the MoE module. The labels 421, 821, and 841 denote different expert-routing settings, where the first number indicates the number of routed experts, the second number indicates the number of selected experts per token, and the last number indicates the use of one shared expert. (a) Rosetta interface energy ΔΔG. (b) Cα RMSD of the six designed CDR loops. (c) Amino acid recovery rate (AAR) of the six designed CDR loops.

**Figure S8.**
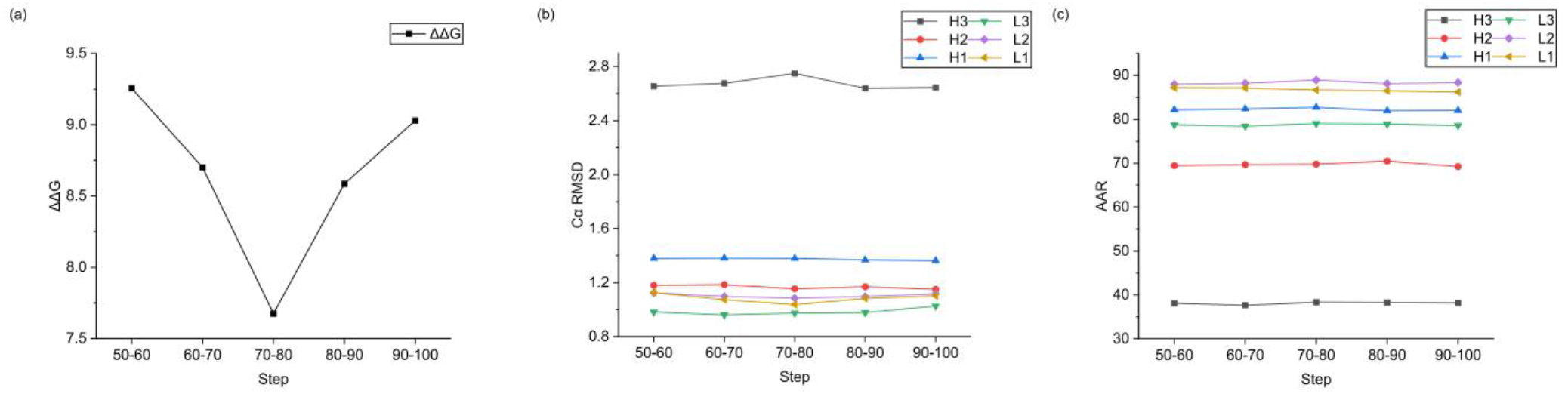
Analysis of the force-guidance schedule under the simultaneous six-CDR design setting. Force-guided sampling was applied over different sampling-step intervals, and the resulting designs were evaluated by interface energetics, structural accuracy, and sequence recovery. (a) Rosetta interface energy ΔΔG. (b) Cα RMSD of the six designed CDR loops. (c) Amino acid recovery rate (AAR) of the six designed CDR loops.

### Case study

We further provide a practical design pipeline for applying ForceFlowAb when an experimentally resolved antibody–antigen complex is unavailable. In this setting, ForceFlowAb was combined with a predefined antibody or nanobody framework and an antigen structure to perform antigen-conditioned CDR-H3 design. As a case study, we used a pre-existing humanized nanobody framework as the starting scaffold. For each antigen, candidate antibody–antigen complex templates were generated by positioning the framework relative to the antigen. The RFdiffusion-based (Bennett et al., 2026) and HDOCK-based (Yan et al., 2017) strategies were applied independently, and each generated 1,000 candidate complex poses. These poses served solely as structural contexts for the subsequent ForceFlowAb design stage and were not treated as final designs. ForceFlowAb then generated one CDR-H3 sequence–structure design from each input pose. The outputs from the two template-generation strategies were retained as separate groups rather than pooled into a single design set.

We compared RFantibody (Bennett et al., 2026) with two ForceFlowAb-based pipelines using RFdiffusion-derived and HDOCK-derived input templates, respectively. All generated candidates were subsequently evaluated using Rosetta interface energy as a computational measure of predicted interface favourability. These results provide an initial assessment of whether ForceFlowAb can be integrated with external template-generation methods to support antigen-conditioned CDR design when no experimentally resolved antibody–antigen complex is available.

**Figure S9.**
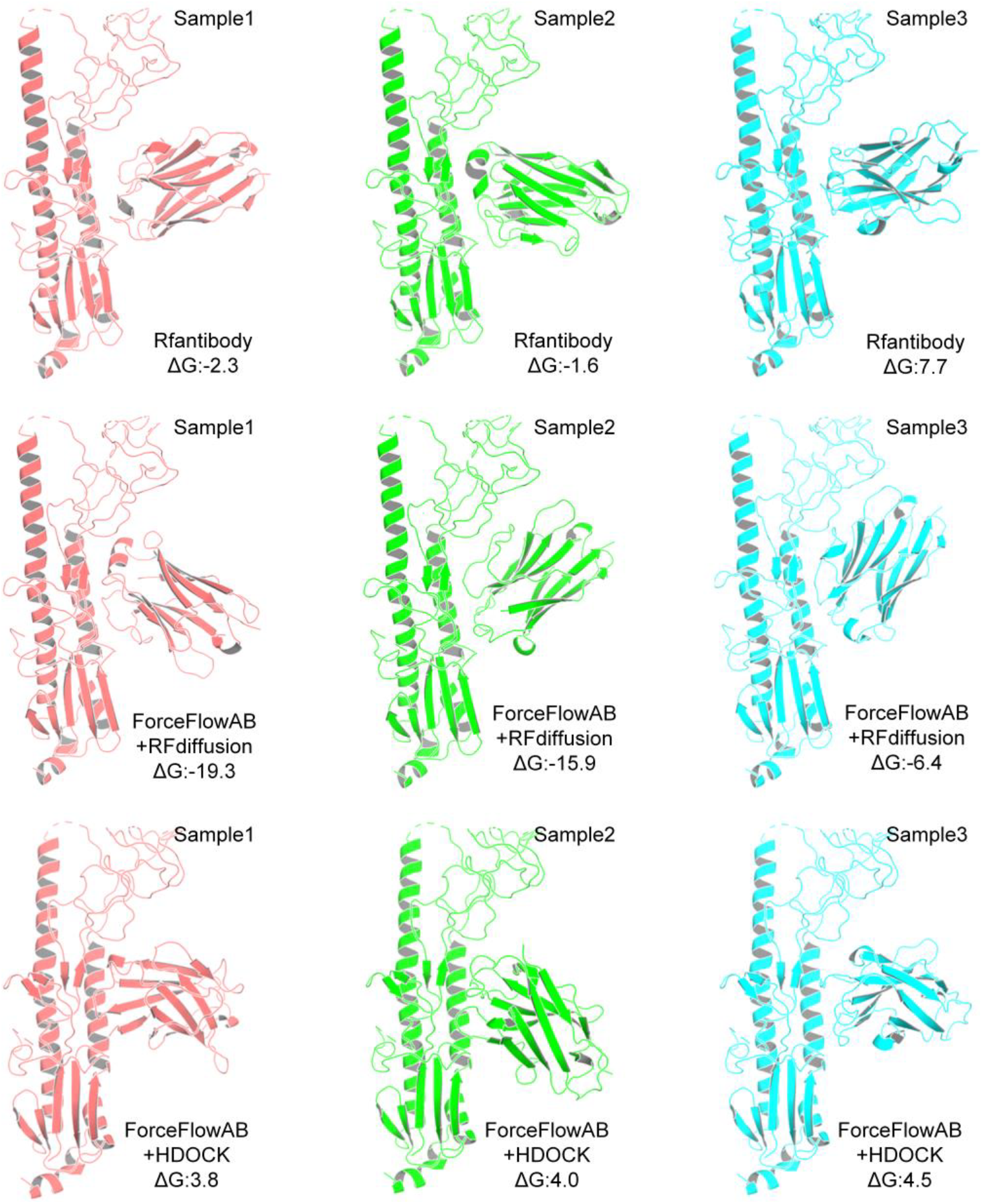
Representative case-study results on flu_HA using the h-NbBCII10 nanobody framework. RFantibody, ForceFlowAb with RFdiffusion-derived templates, and ForceFlowAb with HDOCK-derived templates were compared. Rosetta interface energies (ΔG) are reported for each generated complex.

## Supplementary Note 1

### Model Training

The model used 128-dimensional residue features, 64-dimensional pair features, a six-layer CDR condition inference module consisting of IPA and Transformer encoder blocks, 100 flow steps, and a six-layer interface MoE module with four routed experts, one shared expert, and top-2 expert routing for each residue. Training was performed for 300,000 iterations with a batch size of 16 using the Adam optimizer with β1 = 0.9, β2 = 0.999, an initial learning rate of 1 × 10−4, and weight decay set to 0. Gradients were clipped with a maximum norm of 100. The learning rate was reduced by a factor of 0.8 when the validation metric plateaued, with a minimum learning rate of 5 × 10−6, and an exponential moving average of model parameters was maintained with a decay of 0.995. The same model architecture was used throughout training. During the first 200,000 iterations, the quaternion loss was weighted with α = 1 and the rotation-matrix orientation loss was disabled with β = 0. During the subsequent 100,000 iterations, the loss weights were switched to α = 0 and β = 1, allowing the model to further optimize orientation consistency under the same architecture and optimization settings. Training took approximately three days on a single NVIDIA A100 GPU.

#### Algorithm S1.

Training

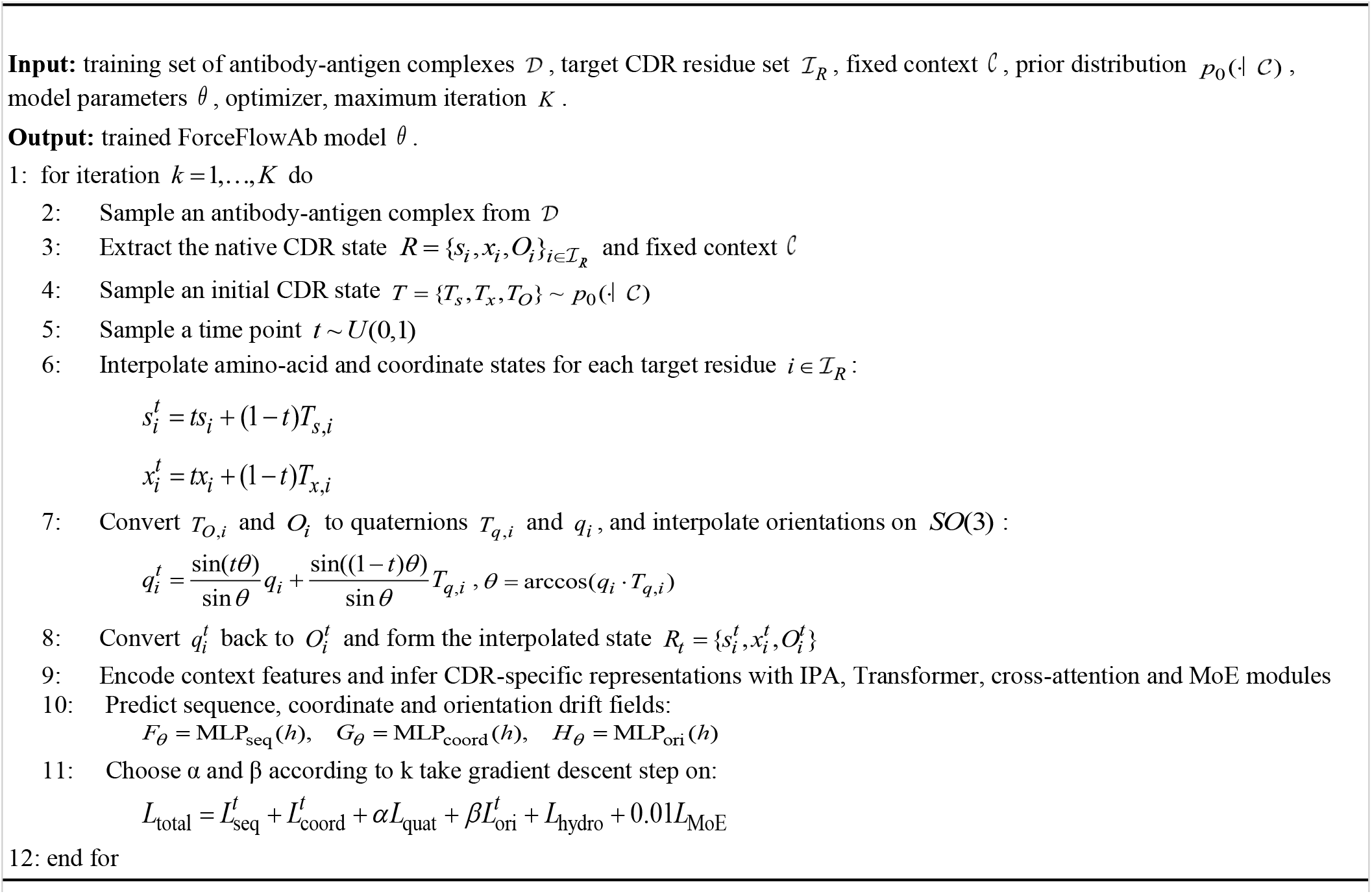

#### Algorithm S2.

Force-guided inference

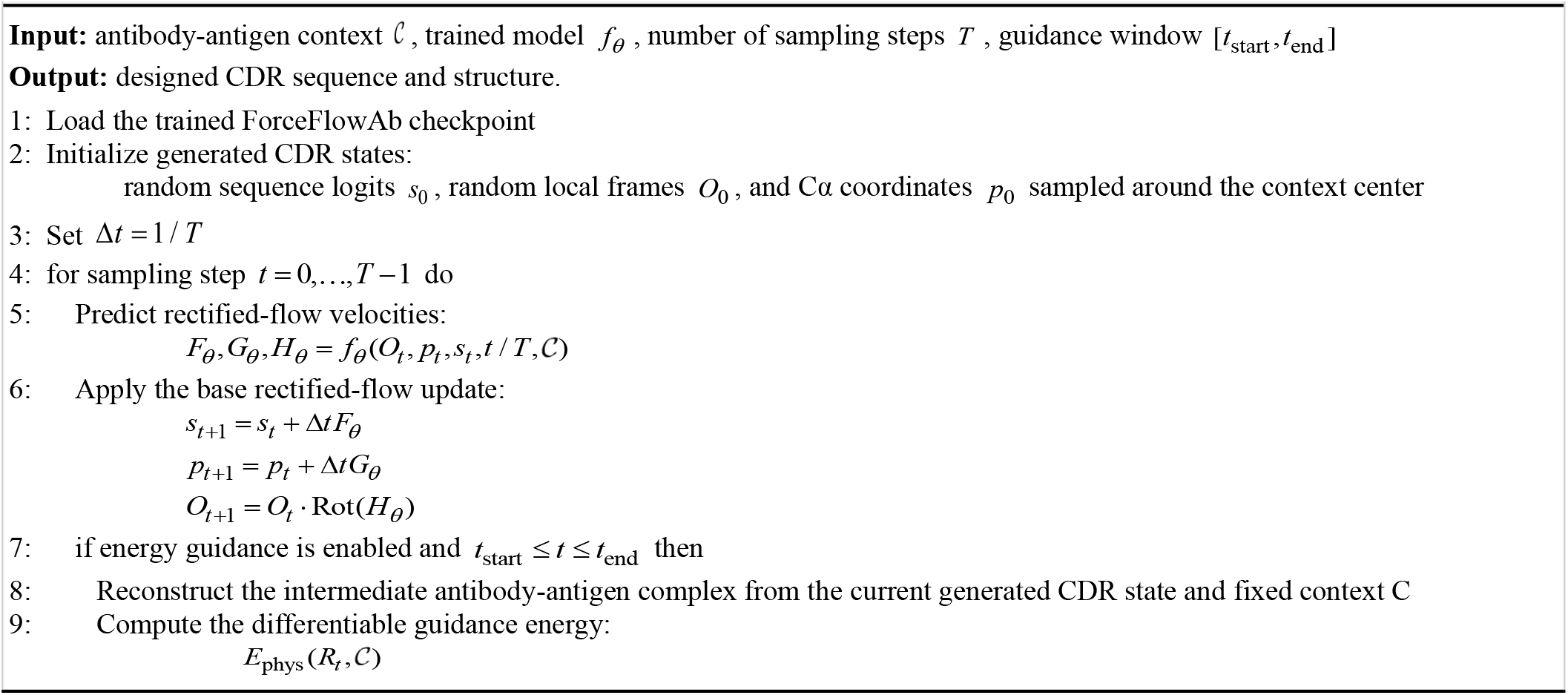
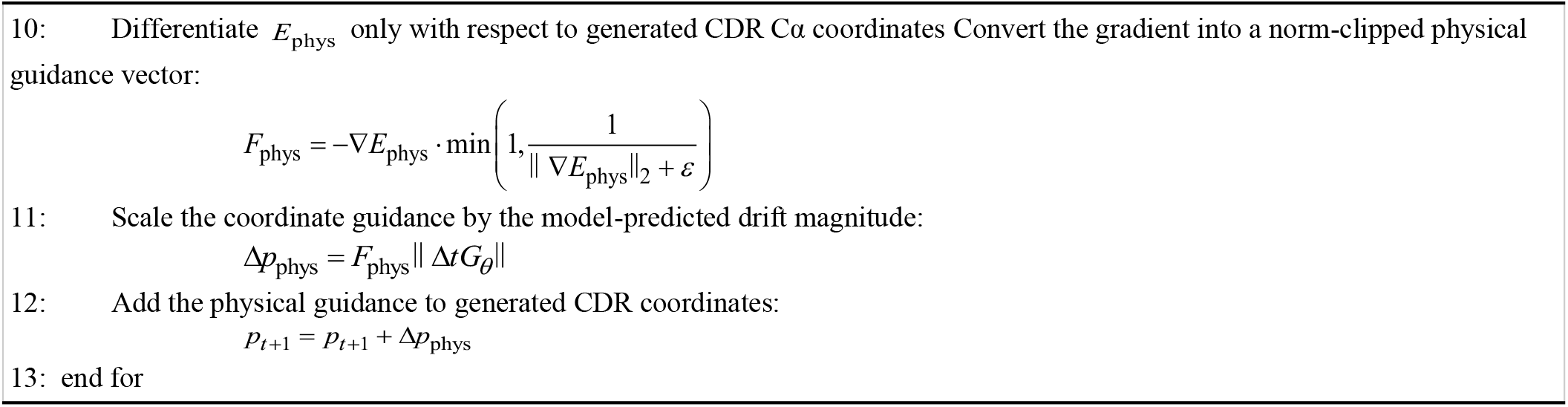

